# Computational Luminance Constancy from Naturalistic Images

**DOI:** 10.1101/358671

**Authors:** Vijay Singh, Nicolas P. Cottaris, Benjamin S. Heasly, David H. Brainard, Johannes Burge

## Abstract

The human visual system supports stable percepts of object color even though the light that reflects from object surfaces varies significantly with the scene illumination. To understand the computations that support stable color perception, we study how estimating a target object’s luminous reflectance factor (LRF; a measure of the light reflected from the object under a standard illuminant) depends on variation in key properties of naturalistic scenes. Specifically, we study how variation in target object reflectance, illumination spectra, and the reflectance of back-ground objects in a scene impact estimation of a target object’s LRF. To do this, we applied supervised statistical learning methods to the simulated excitations of human cone photoreceptors, obtained from labeled naturalistic images. The naturalistic images were rendered with computer graphics. The illumination spectra of the light sources and the reflectance spectra of the surfaces in the scene were generated using statistical models of natural spectral variation. Optimally decoding target object LRF from the responses of a small learned set of task-specific linear receptive fields that operate on a contrast representation of the cone excitations yields estimates that are within 13% of the correct LRF. Our work provides a framework for evaluating how different sources of scene variability limit performance on luminance constancy.

## Introduction

The perceived color of an object has important behavioral implications because color helps to identify objects and their surface properties (Mollon, 1989; Jacobs, 1981). The computational challenge underlying object color perception is that the light reflected from an object depends not just on the object’s surface reflectance, but also on object-extrinsic factors such as the scene illumination, the pose of the object, and the viewpoint of the observer in the scene (Fig. 1a). To compute a perceptual representation of object color that is correlated with the object’s physical surface reflectance, the visual system must account for these object-extrinsic factors. The ability of a visual system to compute a representation of object color that is stable against variation in object-extrinsic factors is called color constancy. A well-studied special case of color constancy is when the stimuli are restricted to be achromatic. This special case is called lightness constancy (Gilchrist, 2006). Although human lightness and color constancy are not perfect, they are often very good (Foster, 2011; Brainard & Radonjić, 2014; Adelson, 2000; Kingdom, 2011).

**Figure 1:**
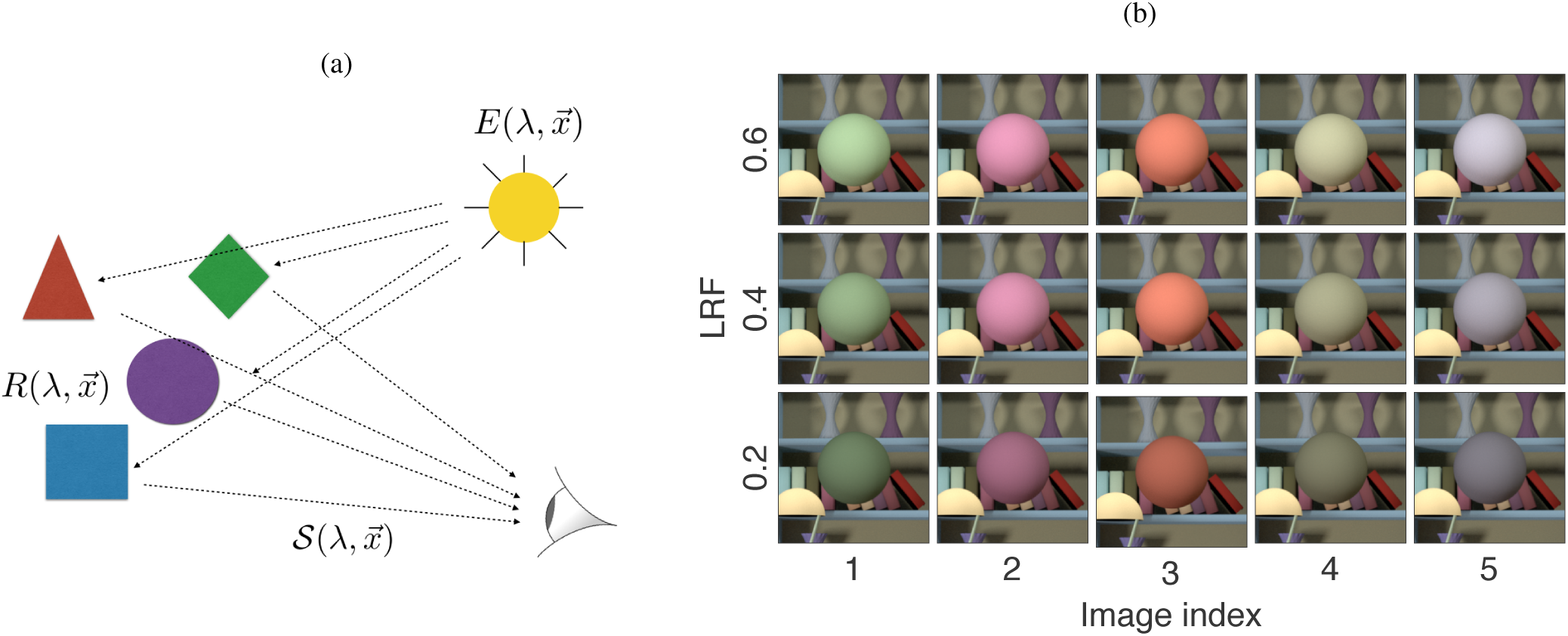
Color and luminance constancy: (a) The light reflected from an object to the eye depends both on the surface reflectance of the object and on the scene illumination. The reflected light also depends on geometric factors, such as the object’s shape, pose, and position. The human visual system accounts for variations in reflected light due to object-extrinsic factors and produces a percept of object color that is relatively stable across such variations. (b) Images of a sphere under a fixed illuminant. Down each column, the sphere’s luminous reflectance factor (LRF) varies, but its relative reflectance spectrum (i.e. its spectral shape) is held fixed. Across each row, the relative reflectance spectrum varies, but the LRF is held fixed. We cast the problem of computational luminance constancy to be that of estimating the LRF of a target object from the image, across variation in other scene factors. Specifically, we study variation in target object relative surface reflectance, variation in illumination spectrum, and variation in the surface reflectance of the other objects in the scene.

**Figure 2:**
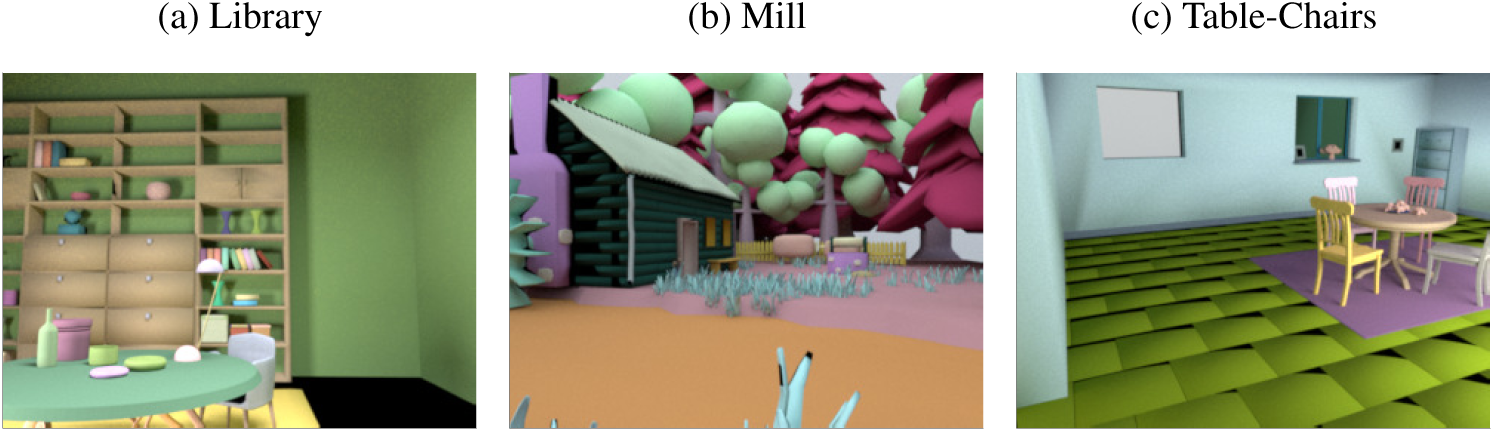
Base scenes. Each image shows a rendering of one base scene without additional inserted objects. The reflectance spectra of the distinct surfaces and the spectral power distributions of the light sources in each scene have been assigned randomly from statistical models of naturally occurring spectra. The images have been scaled and tone mapped individually for illustrative purposes.

The computational problem of color constancy may be framed as how to obtain stable descriptions of the spectral surface reflectance functions of the objects in a scene. Early work on computational color constancy considered a simplified imaging model where scenes consisted of multiple flat matte objects and a single spatially diffuse illuminant (Land, 1977; Buchsbaum, 1980; Maloney & Wandell, 1986). Subsequent work incorporated probabilistic descriptions of the statistics of naturally occurring scenes (D’Zmura & Iverson, 1994; D’Zmura, Iverson, & Singer, 1995; Brainard & Freeman, 1997). A key insight from these computational studies is that stable color descriptors of a target object cannot be obtained solely from the light reflected from that target object. It is possible, however, to obtain stable color descriptors by jointly analyzing the light reflected from multiple objects in the scene. As a consequence, color constancy will be affected both by variation in the illumination and by variation in the surface reflectance of other objects in the scene. Color constancy algorithms should therefore be evaluated with respect to variation in both of these factors (Brainard & Wandell, 1986; Brainard & Freeman, 1997).

Recently, supervised statistical learning methods have been applied to the computational problem of color constancy. These methods use the joint statistics of labeled images and scene variables to learn mappings that extract stable surface color descriptors from image input (Barron, 2015). This general approach has been useful for other perceptual problems in early and mid-level vision. For example, a recently developed technique called accuracy maximization analysis (AMA) learns linear receptive fields optimized for particular perceptual tasks (Geisler, Najemnik, & Ing, 2009; Burge & Jaini, 2017; Jaini & Burge, 2017). AMA has been used to develop ideal observers for estimating defocus blur, binocular disparity, and retinal motion estimation with natural stimuli. These ideal observers provide excellent predictions of human performance (Burge & Geisler, 2011, 2012, 2014, 2015).

Supervised learning requires large labeled datasets. Such datasets are not readily available for the study of color constancy. Although there are datasets of calibrated color images, these do not provide ground truth information about surface reflectance and illumination at each image location (Chakrabarti & Zickler, 2011; Nascimento, Amano, & Foster, 2016; Parraga, Brelstaff, Troscianko, & Moorehead, 1998; Tkacik, G., Garrigan, P., Ratliff, C., Milcinski, G., Klein, J. M., Sterling, P., Brainard, D. H., & Balasubramanian, V., 2011; Skauli & Farrell, 2013; Olmos & Kingdom, 2004). There are a few databases consisting of images of posed scenes where surface reflectances of individual objects have been measured (Funt & Drew, 1988; Ciurea & Funt, 2003), but these databases are not large enough to drive supervised learning. Finally, recent work has applied crowd-sourcing techniques to annotate natural images with surface reflectance descriptors (Bell, Bala, & Snavely, 2014).

In this paper, we use high-quality computer graphics to generate large datasets of naturalistic images where the surface reflectance corresponding to each image pixel is known. This approach allows us to investigate computational color constancy with naturalistic stimuli, while retaining the ability to control the properties of objects and illuminants. We tackle the specific problem of luminance constancy, a constitutive component of the more general color constancy problem (Fig. 1b).

We define the computational problem of luminance constancy^1^ as that of estimating the luminous reflectance factor (LRF) of a target object’s surface reflectance function. The LRF is a measure of the overall amount of light reflected by a surface relative to the reference illuminant itself^2^ (American Society for Testing and Materials, 2017). Obtaining the LRF from a known surface reflectance function requires two steps. First, one computes the luminance of the light that would be reflected from the surface under a reference illuminant. Second, one normalizes the result by the luminance of the reference illuminant itself. Here we use CIE daylight D65 as the reference illuminant and the CIE 1931 photopic luminosity function to compute luminance (Commission Internationale de l’éclairage, 1986). Sometimes LRFs are expressed as a percentage. Here we express LRFs as proportions, so that they take on values between 0 and 1. An LRF of 0 means that none of the light from the reference illuminant is reflected from the surface. An LRF of 1 means that the luminance of the surface reflectance under the reference illuminant is the same as the luminance of a perfect reflector under that same illuminant.

## Methods

### Overview

There are four key steps to our methods. First, we generate a labeled set of training images. Second, we use a model of the early visual system to compute the responses of the cone photoreceptor mosaic to the labeled images. Third, we learn the receptive fields (RFs) that extract the most useful information from the cone responses for the task. Fourth, we evaluate how well the responses of the receptive fields may be decoded to achieve luminance constancy.

### Labeled training data

#### Virtual naturalistic scenes

The light that reflects from objects to the eyes depends on many factors. These include the surface reflectance, texture, material and geometry of the objects, the spectral power distribution of the illuminants, and the position of the observer. We have developed a rendering package (https://github.com/BrainardLab/VirtualWorldColorConstancy) that allows us to construct models of naturalistic scenes, with key scene factors under programmatic control. The package builds on the RenderToolbox4 package (http://rendertoolbox.org; Heasly, Cottaris, Lichtman, Xiao, & Brainard, 2014), and harnesses the open-source computer graphics renderer Mitsuba (https://www.mitsuba-renderer.org, Jakob, 2010) to produce physically-accurate images from the scene models. Because the images are rendered from known scene models, each image pixel can be labeled with the surface reflectance of the corresponding object. By using statistical models of daylight illumination and object surface reflectance to guide the rendering of scenes, the package allows us to produce large labeled sets of images that capture key aspects of the task-relevant statistical structure of natural scenes.

The RenderToolbox4 package includes a collection of base scenes (Fig. 2). Base scenes specify an arrangement of objects and light sources. Base scenes may be enriched by the insertion of additional objects, chosen from an object library (Fig. 3), and by the insertion of additional light sources. Once the position, size, and pose of the inserted objects and light sources have been set, a surface reflectance function can be assigned to each object in the scene, and a spectral power distribution function can be assigned to each light source (Fig. 4). The resulting scene model can then be rendered from any specified viewpoint.

**Figure 3:**
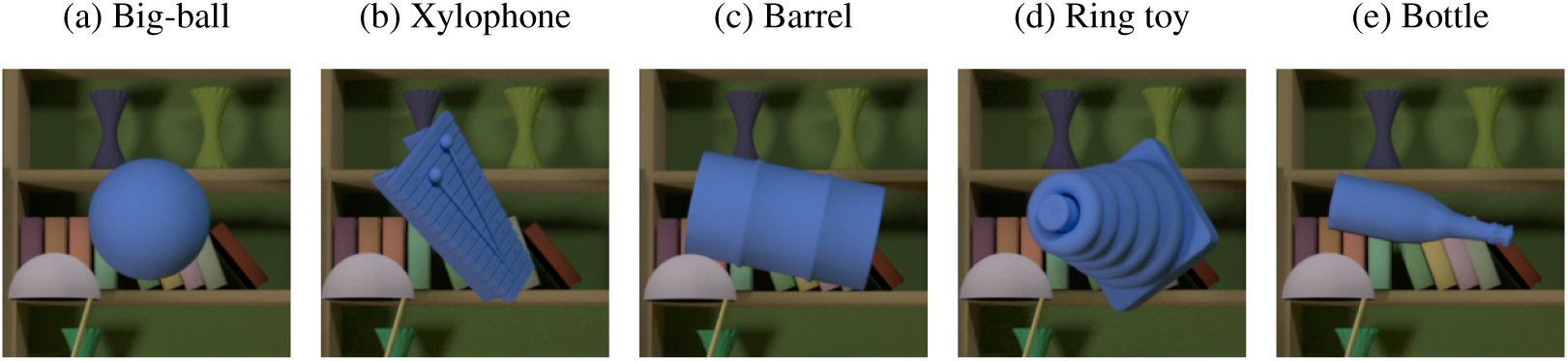
Library base scene with inserted objects. The rendering package can be used to insert objects into base scenes. Each image shows a different object inserted into the Library base scene. The rendering viewpoint was set so that the object was at the center of the images. The full rendered images have been cropped so that the inserted objects are easily visible.

**Figure 4:**
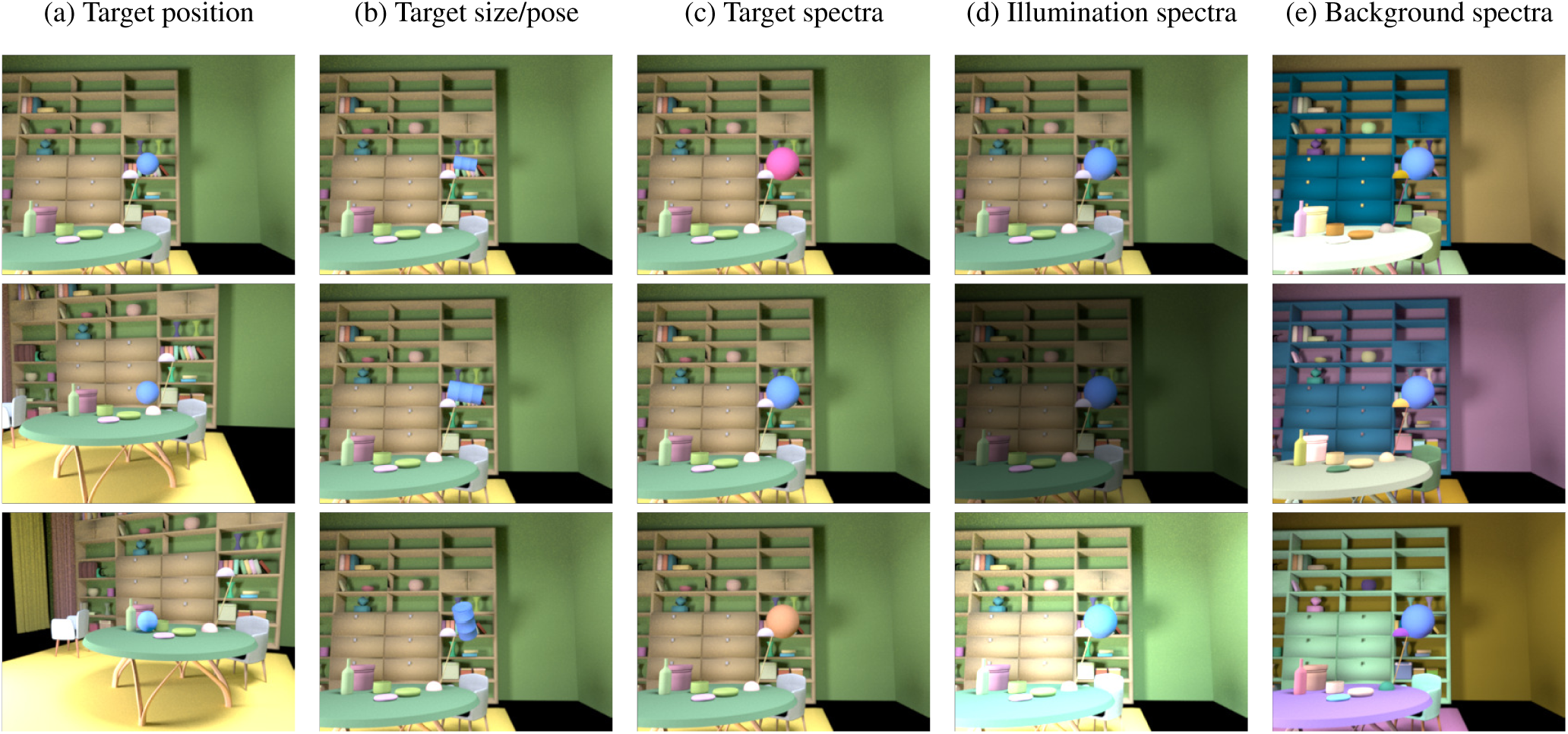
Scene transformations. The properties of a visual scene can be broadly classified into two groups: geometrical (a-b) and spectral (c-e). Our rendering package provides control over these properties as illustrated by the columns of the figure. (a) Variation in inserted object positions. (b) Variation in object size and pose. (c) Variation in the surface reflectance of the target object. (d) Variation in the spectral power distributions of the light sources. (e) Variation in surface reflectance of the background objects. All images in this figure are rendered with common scaling and tone mapping.

In the present work, we used our package to generate datasets of naturalistic scenes and corresponding images. We used one fixed base scene and inserted a spherical target object of fixed size into this scene. All surfaces in the scene model were matte and did not have specularities. There were multiple light sources in the scene, and the rendering process simulated shadows as well as mutual reflection of light between nearby surfaces. We generated three distinct datasets, which we refer to as Conditions 1-3 (Fig. 5). These are described below. Across these datasets the luminance constancy problem (i.e. estimating the target object LRF) becomes progressively more difficult.

**Figure 5:**
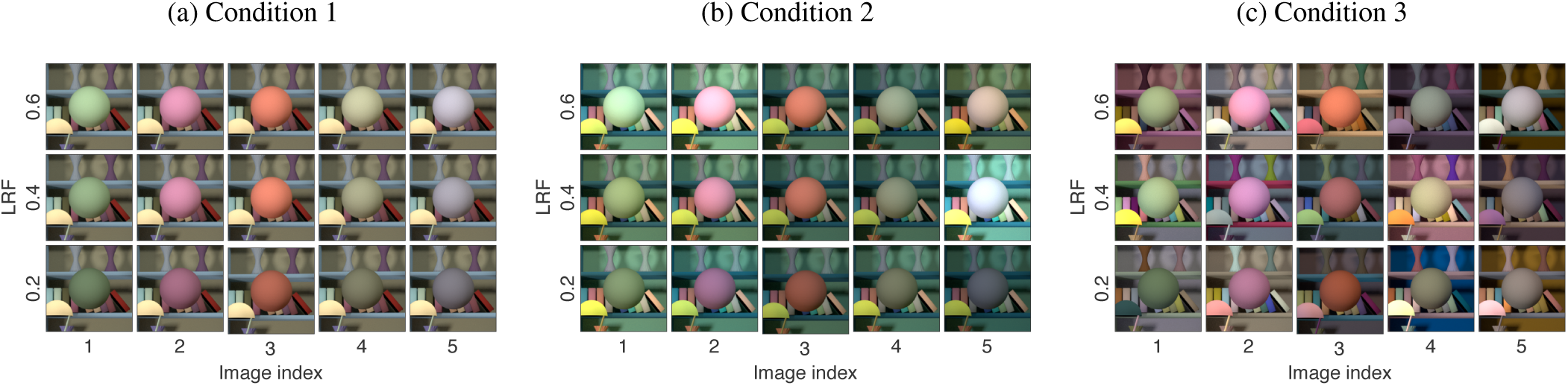
Conditions 1-3: Illustrative images for the three types of spectral variations studied in this paper. (a) Condition 1: variable target object relative reflectance spectrum, fixed light source spectra, fixed background object spectra (same as Fig. 1b). (b). Condition 2: variable target object relative reflectance spectrum, variable light source spectra, fixed background object spectra.(c) Condition 3: variable target object relative reflectance spectrum, variable light source spectra, variable background object spectra. The spheres in each row of each panel have the same LRF, while the spheres in each column of each panel have the same relative surface reflectance spectrum. Across panels a-c, spheres in corresponding locations have the same surface reflectance. In all three panels, the overall light source spectra scale factors (see Methods) were drawn from a uniform distribution on the range [0.5, 1.5]. This is smaller than the variation we studied, but allows us to show the variation within the dynamic range available for display in publication. All images in this figure are rendered with common scaling and tone mapping.

The variation within our datasets captures the essence of the computational problem of lightness constancy up to effects of scene geometry, an additional richness that we do not address in this paper (see Funt & Drew, 1988; Barron & Malik, 2012).

In Condition 1, the relative reflectance spectrum of the target object was the only source of scene variation other than the target object LRF. The reflectance spectra of the background objects and the spectral power distribution of the light sources were held fixed. In Condition 2, the spectral power distribution of the light sources varied in addition to the relative reflectance spectrum of the target object. Finally, in Condition 3 three distinct scene factors–target object relative reflectance spectrum, spectral power distribution of the light sources, and reflectance spectra of the background objects –varied in addition to the target object LRF. The datasets for Conditions 1 and 2 consisted of 1000 scenes and images, 100 for each of 10 target object LRF values. The dataset for Condition 3 consisted of 3000 scenes and images, 300 each for the 10 target LRF values. The LRF values were equally spaced between 0.2 and 0.6. More than 90% of the surface reflectance spectra (generated as described below) fell within this range. For each LRF value, we generated a different relative target object surface reflectance for each scene.

We used the Library base scene and a spherical target object. The Library base scene contains 2 area lights. We inserted one additional spherical light source into the scene. In Conditions 2 and 3, the overall intensities of the three light source illumination spectra were equal, while their relative shape varied. The overall intensity varied from scene to scene. The position and size of the inserted object, the inserted light source, and the viewpoint on the scene were held fixed across all scenes. The geometry is shown in Figure 9a below. Surface and illuminant spectra were sampled according to statistical models of naturally occurring spectra. Multispectral images (320 × 240 pixels) were rendered at 31 evenly-spaced wavelengths between 400nm and 700nm.

#### Illumination spectra

To generate illumination spectra, we developed a statistical model of the Granada daylight measurements (Hernández-Andrés, Romero, Nieves, & Lee, 2001; http://colorimaginglab.ugr.es/pages/Data) and then sampled randomly from the model. To match the wavelength spacing we use in rendering, we resampled the wavelength spacing of the Granada spectra to 31 evenly-spaced wavelengths between 400 nm and 700 nm.

Our statistical model approximates normalized spectra from the Granada dataset using the first 6 principle components of the normalized measurements from that dataset. The distribution of weights on these components is approximated with a 6- dimensional Gaussian, where the mean and covariance matrix match the sample mean and covariance of the weights (Brainard & Freeman, 1997). To generate a random relative illuminant spectrum, we sample weights randomly from this Gaussian and then perform a weighted sum of the components. Figure 6b illustrates the relative spectra of draws obtained using this procedure. Chromaticities and rendered color renditions of these draws are shown in Figures 6c and 6d. Details of the statistical model for illumination are provided in the appendix.

**Figure 6:**
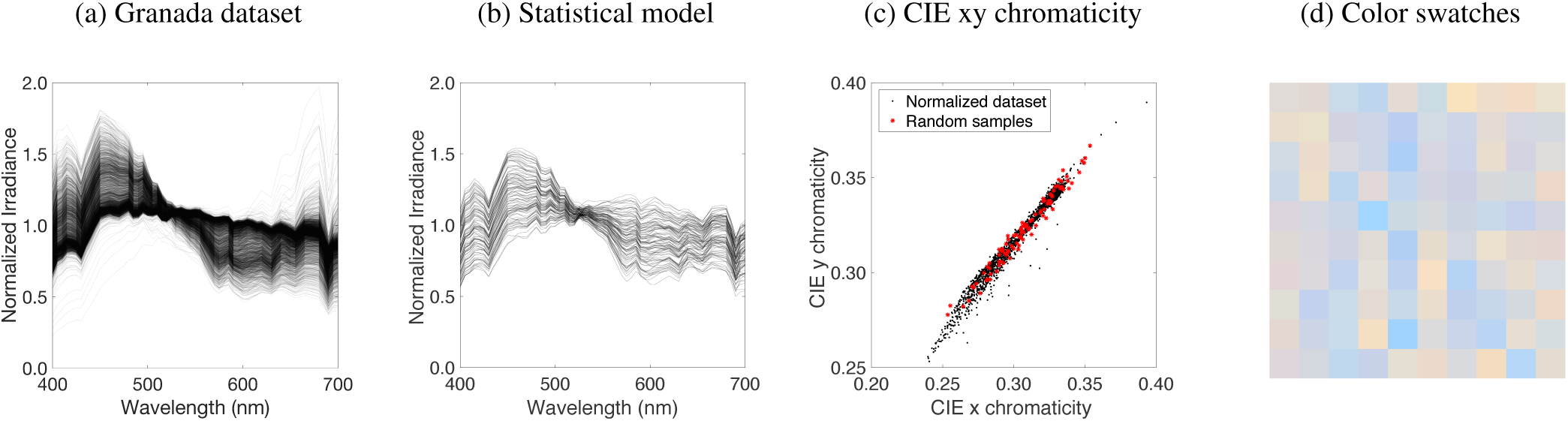
Statistical model of illumination spectra: (a) Normalized Granada dataset. Each spectrum is normalized by its mean power. (b) Sample spectra generated using the statistical model derived from normalized Granada dataset. (c) CIE xy chromaticities of the Granda dataset (black) and the samples shown in panel b (red). (d) sRGB renditions of the samples shown in panel b.

The multivariate Gaussian model described above is based on normalized spectra. Thus, it does not embody the large variation in overall intensity of natural daylights. To restore the overall intensity variation in the Granada dataset, which spans approximately three orders of magnitude, we scaled each randomly generated spectrum with a random number sampled from a log-uniform distribution spanning three orders of magnitude. The particular range of scale factors was chosen so that the mean number of cone isomerizations from the target object ranged from approximately 100 to 100,000, given a 100 ms cone integration time (see below). This range of cone isomerizations corresponds approximately to that of natural daylight viewing.

#### Surface reflectance spectra

To generate random reflectance spectra, we employed principles similar to those used to generate random illuminant spectral power distributions. We used measurements from the Munsell (Kelly, Gibson, & Nickerson, 1943) and Vrhel (Vrhel, Gershon, & Iwan, 1994) surface reflectance datasets to create a dataset containing 632 reflectance spectra (462 from the Munsell data and 170 from the Vrhel data; Figure 7a). We resampled the spectra to 31 evenly-spaced wavelengths between 400 nm and 700 nm. Figures 7b, 7c, and 7d illustrate reflectance sampled obtained using the model. Details of the statistical model for surface reflectance spectra are provided in the appendix.

**Figure 7:**
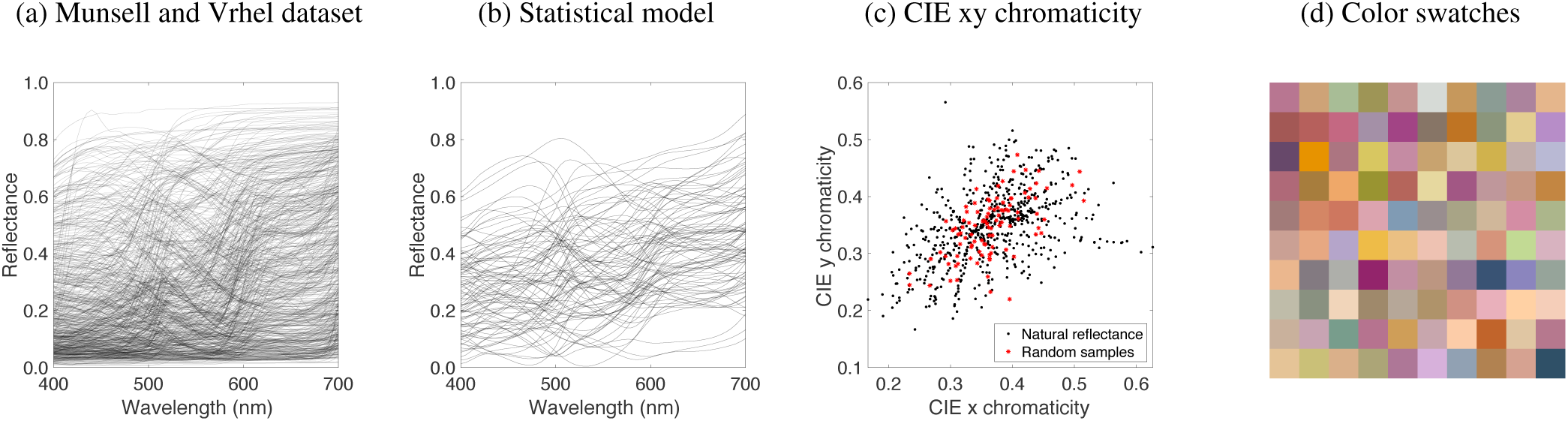
Statistical model of surface reflectance: (a) Surface reflectance spectra from the Munsell and Vrhel datasets. (b) Sample spectra generated using the surface reflectance statistical model. (c) CIE xy chromaticity diagram of the Munsell and Vrhel reflectances (black) and the sample spectra shown in panel b (red). (d) sRGB renditions of the samples shown in panel b, rendered under the CIE D65 daylight spectrum.

To generate the target object reflectance at a particular LRF, the sampled reflectance spectrum was scaled such that its LRF had the desired value (see appendix). Figure 8 shows color renderings of target reflectance spectra under CIE illuminant D65, for the evenly spaced LRF values we studied.

**Figure 8:**
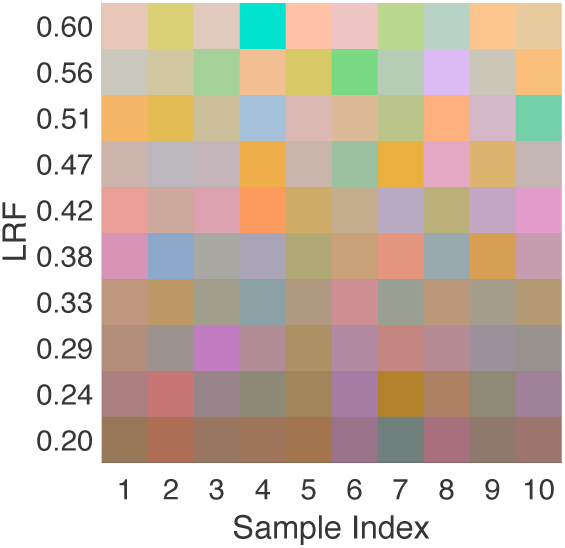
Target object surface reflectance spectra: sRGB renditions of the target object surface reflectance rendered under CIE D65 daylight spectrum. The figure shows 10 random samples at 10 equally spaced LRF levels in the range [0.2, 0.6]. Each row contains 10 random samples of reflectance spectra generated at the LRF shown on the left.

**Figure 9:**
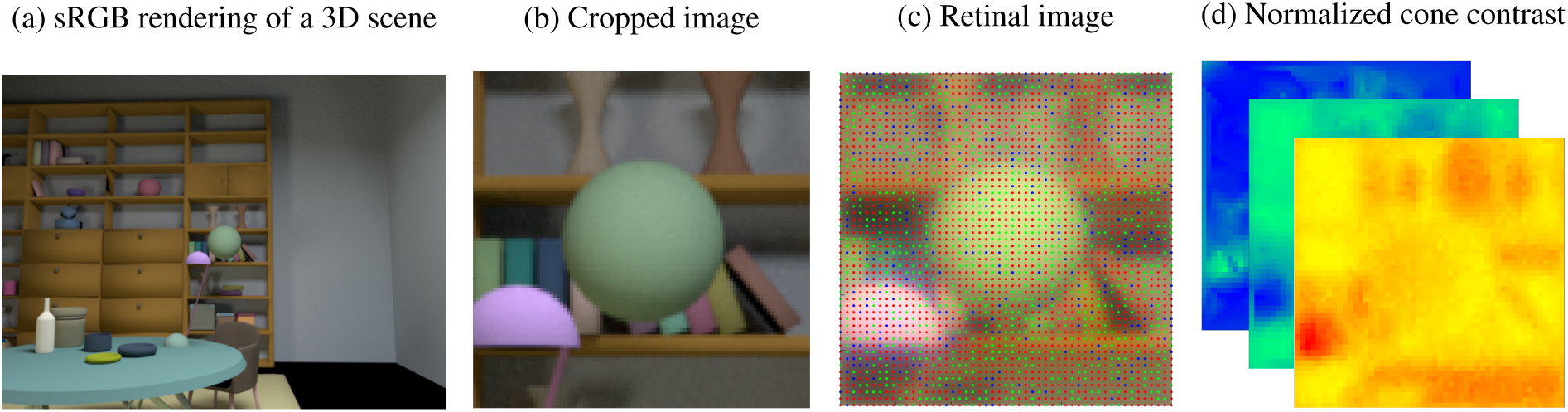
Pipeline for generating labeled datasets: The labeled images used in this paper are generated as follows: (a) A 3D virtual scene, containing a target object is created. The rendering viewpoint is selected so that the target object is in the center of the image. Spectra are assigned to the target object, illuminants, and other objects in the scene using the statistical models described in the text. A multispectral image of the scene is then rendered. (b) The central portion of the rendered image is cropped around the target object. (c) The retinal image corresponding to the rendered image is computed, which is then used to compute the number of photopigment isomerizations in the cone photoreceptors. The figure shows a rendering of the retinal image after optical blurring, with the location and identity of the cones indicated by an overlay (L cones: red, M cones: green, S cones: blue). (d) The cone excitations are linearly interpolated to estimate the responses of all the three types of cone at each location (demosaicing). Finally, the demosaiced images are contrast normalized.

### Model of early visual system

Light entering the eye is focussed and blurred by eye’s optics to form the retinal image. This image is then sampled by a mosaic of cone photoreceptors. The excitations of these photoreceptors provide the information available to the neural visual system for further processing. Because we are interested in how well luminance constancy may be achieved by the human visual system, we simulated the cone excitations to our scenes using a model of the early visual system.

We focused our analysis on image regions local to the target by cropping the rendered images to 1° ×1° degrees of visual angle around the target object (51 × 51 pixels; Figure 9b). The local analysis is motivated by the fact that neural receptive fields early in the visual pathways (e.g., retina, primary visual cortex) pool information locally. In primary visual cortex, foveal receptive fields have a maximum spatial extent of approximately 1 degree of visual angle (Gattass, Gross, & Sandell, 1981; Gattass, Sousa, & Gross, 1988). We sought to understand how well responses from AMA-learned receptive fields at a similar scale could be used to achieve luminance constancy.

We modeled a visual system having typical optical blurring (including axial chromatic aberration, Marimont & Wandell, 1994), and typical spatial sampling with an interleaved rectangular mosaic of long (L), middle (M) and short (S) wavelength-sensitive cones (Brainard, 2015; Fig. 9c). The cone mosaic contained L:M:S cones in approximately the ratio 0.6:0.3:0.1 (1523 L cones, 801 M cones, 277 S cones) and with spectral sensitivities derived from the CIE physiological standard (Commission Internationale de l’éclairage, 1986). Cone excitations were taken as the number of photopigment isomerizations, assuming an integration time of 100 msec. The Poisson nature of the photopigment isomerization was also included (Hecht, Shlaer, & Pirenne, 1942). This modeling was implemented using the software infrastructure provided by ISETBio (https://isetbio.org).

To put the computed cone excitations into a form for further analysis, we demosaiced the excitations for each cone class using linear interpolation to obtain a cone excitation image for each cone class. The lens and macular pigment absorb more short than long wavelength light, so that the excitations of the S cones tend to be smaller than those of the L and M cones. To make the magnitude of the three cone excitation images similar across cone classes, we normalized each cone excitation image by the summed (over wavelength) quantal efficiency of the corresponding cone class. The demosaicing and normalization do not alter the available information relative to that carried by the cone excitations themselves.

To capture key properties of post-receptoral processing, such as contrast normalization (Heeger, 1992; Albrecht & Geisler, 1991; Carandini & Heeger, 2012), we processed the cone excitation images as follows: We first converted each cone excitation image to a corresponding cone contrast image. This was accomplished by computing the mean response over the three cone excitation images, and then subtracting off and dividing by this mean. To model contrast normalization, we then divided the contrast images by the sum of squared contrasts taken over image pixels and cone classes.

### Computational luminance constancy

We used our datasets to determine how well target object LRF can be estimated from cone excitations and from normalized cone contrasts. Studying both representations allows us to understand how early contrast coding and normalization affect luminance constancy. We applied accuracy maximization analysis (AMA) to learn the optimal receptive fields for estimating LRF, and evaluated performance when the responses of these receptive fields are optimally decoded.

#### Learning optimal receptive fields

AMA is a task-specific Bayesian method for dimensionality reduction. When provided with a labeled training set, a receptive field response model, a decoder that uses these responses to estimate the stimulus label, and a cost function, AMA returns a set of N linear receptive fields, where N is a parameter of the procedure. These receptive fields are chosen so that decoding their responses leads to minimum expected-cost estimation, relative to any other choice of N linear receptive fields. This is possible because given a training set together with a set of N linear receptive fields and an explicit cost function, it is possible to determine the minimum expected-cost estimator that maps receptive field responses to the stimulus labels. Moreover, a close approximation to the corresponding expected estimation cost may be explicitly computed. Thus, AMA searches over the space of linear receptive fields to find the set that minimizes the expected estimation cost. For each condition, we learned the receptive fields with a training set consisting of 90% of the images.

In our implementation of AMA, we used both the Kullback-Leibler divergence cost function (corresponding to the maximum a posteriori estimator) and the mean squared error cost function (corresponding to the posterior mean estimator) to evaluate the accuracy of the AMA estimates of LRF. Training with both cost functions yielded similar estimation performance; the results reported here are for when the Kullback-Leibler divergence cost function was used in training. We chose this cost function because the estimates it yields at the boundaries of the LRF range are less biased than those obtained with the mean squared error cost function.

Details of how AMA learns receptive fields and how the receptive field responses are optimally decoded are provided in previously published work (Geisler et al., 2009; Burge & Jaini, 2017; Jaini & Burge, 2017).

#### Decoding optimal receptive fields

Once the AMA receptive fields have been learned, a general decoder is needed that can be used with arbitrary test images. A general decoder is necessary because the decoder used to learn the receptive fields can be used only with the training set. Specifically, the decoder used to learn the receptive fields requires the response mean and response variance of each receptive field to every labelled stimulus in the training set (Geisler et al., 2009; Burge & Jaini, 2017). To proceed, first we use the AMA receptive fields and the training dataset to find the distributions of the receptive field responses conditional on the target object LRFs. Then, we model these conditional response distributions with multivariate Gaussians. To estimate the LRF of the target object in a test image, we use Bayes’ rule together with the Gaussian conditional distributions and a uniform prior to obtain the posterior distribution over LRF values, given the receptive field responses. The optimal estimate is the LRF value that minimizes the expected value of the cost function, where the expectation is taken over the posterior. For the results we report, we again used the Kullback-Leibler divergence cost function.

The Gaussian approximations for the posterior were estimated using the same training set that was used to learn the receptive fields. Estimation performance was evaluated on a test set consisting of the 10% of images in the dataset that were not in the training set. For our primary results, estimation was based on six receptive fields. The AMA analysis package is available at: https://github.com/BrainardLab/LuminanceConstancyAmaAnalysis.

#### Baseline methods

To provide baselines for evaluation of the estimation method described above, we used linear regression. First, we solved for the weights on the average L, M and S cone excitations corresponding to the target object that best predicted the target LRF values. We took the average cone responses from a 3 × 3 pixel region at the center of the target object. This baseline method disregards information carried by the image regions corresponding to the background objects. We also performed regression on the contrast normalized version of the cone excitation images. This second baseline method indirectly incorporates information carried in the image regions corresponding to the background objects, through their effect on the contrast normalized responses at the center of the target object. We also evaluated the performance of a naive model that uses the mean LRF value in the training set as the estimate, irrespective of the image data; that is, the naive model always estimated the LRF to be equal to 0.4.

#### Quantifying estimation performance

We quantified the performance of AMA and the baseline methods at estimating LRF with relative root mean squared error (relative RMSE). Relative RMSE is the square root of the mean of the squared difference between the estimated and true LRF divided by the true LRF. The mean is taken over all stimuli in the test set.

## Results

### Condition 1

In Condition 1, only the LRF and the relative reflectance spectrum of the target object vary across scenes. We used accuracy maximization analysis (AMA) to learn a set of linear receptive fields that are optimal for estimating target LRF from the cone excitations in this condition. Decoding performance is shown in Figure 10a. Figure 10a also shows the performance of the baseline linear regression method. Both methods are trained on 90% of the images in the dataset and tested on the remaining 10%. The LRF estimates obtained for both methods are essentially perfect.

**Figure 10:**
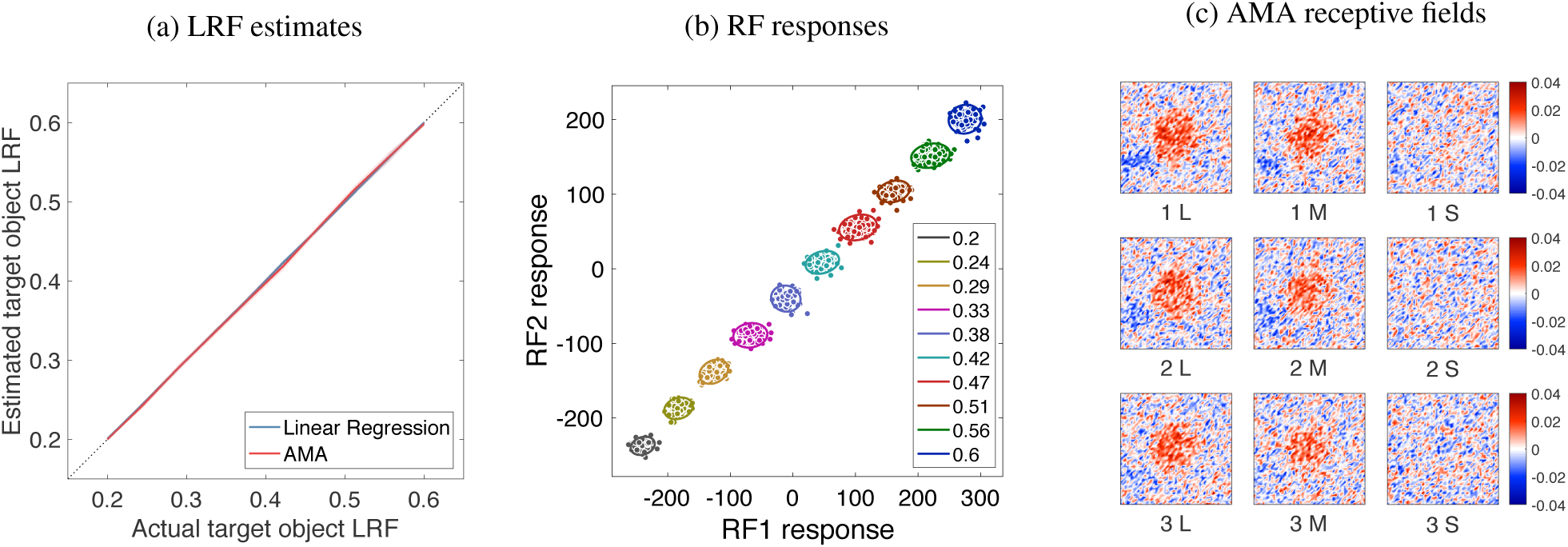
Condition 1 results: (a) Mean estimated LRF ± 1 standard deviation for images in the test set obtained using the AMA-based estimator and linear regression on the cone excitations. Solid lines show the mean estimate, the filled region in lighter color shows ± 1 standard deviation. The diagonal identity line (dotted blue) indicates veridical estimation. The filled regions representing standard deviations are too small to be visible. Relative RMSE (see Fig.14) is 0.7% for linear regression and 1.1% for AMA-based estimation. (b) Training image cone excitations in Condition 1 projected onto the first two AMA receptive fields. Each cluster of responses represents the image patches associated with a particular LRF. The response clusters are approximated by a multivariate Gaussians whose means and covariances are represented by the ellipses shown in the figure.(c) The first three AMA receptive fields. These are specified over the same 1 x 1 degree patch as the stimuli.

Figure 10b shows the responses of the first two AMA receptive fields to all the image patches in the training set. Each individual point represents the receptive field responses to an individual image patch and is color coded by target object LRF. The responses segregate according to the target LRF, which supports the high level of estimation performance shown in Fig-ure 10a.LRF Note that even the responses of the first receptive field separate well (vertical lines would accurately distinguish the LRFs), so that that one receptive field is sufficient for accurately performing the task.

Figure 10c shows the first three AMA receptive fields. Each receptive field performs a weighted sum of the L, M, and S cone excitations. The L and M cone excitations at the target object location receive large weights. The L and M cone excitations at the background object locations and the S cone excitations at all locations receive small weights and are thus largely ignored. For this condition, the background regions provide very little task-relevant information. In addition, luminance is primarily determined by L and M cones, so the S cones should make little direct contribution. The first three receptive fields are similar to each other, reflecting the fact that here the main benefit of multiple receptive fields is to reduce the impact of noise on estimation performance.

The results in Figure 10 were obtained for one particular draw from our statistical model of illumination. We verified that performance remained excellent for other draws.

### Condition 2

Condition 2 includes variation in the spectral power distribution of the light sources, in addition to the variation present in Condition 1. This illumination variation makes the task more difficult because it causes variation in the cone excitations that is not due to target object LRF. We learned AMA receptive fields on the cone excitations to the images in this new condition, and again evaluated decoding performance (Fig. 11(a,b)). Performance is poor. Indeed, the estimates deviate considerably from the true LRF, as seen by the fact that the mean estimate (red line) does not lie on the positive diagonal and by the fact that there is large estimation variability for each value of the true LRF. For this case, linear regression also does poorly. Indeed, the linear regression estimates are essentially the same as would be obtained by simply guessing the mean LRF of the training set (0.4).

**Figure 11:**
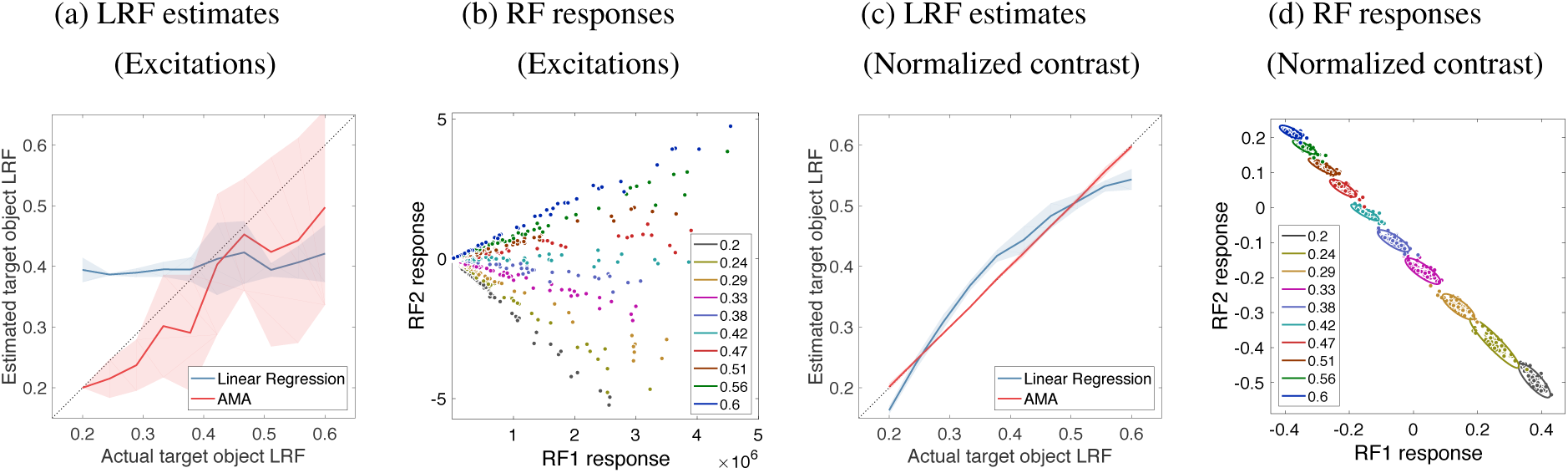
Condition 2 results: (a) Mean estimated LRF ± 1 standard deviation for images in the test set obtained using the AMA-based estimator and linear regression from the cone excitations. The relative RMSE is high (41.4% for linear regression and 26.4% for AMA-based estimation). (b) Cone excitations projected onto AMA receptive fields. (c) Mean estimated LRF ± 1 standard deviation for images in the test set obtained using the AMA-based estimator and linear regression from the contrast normalized cone excitations. The relative RMSE of estimation is 9.3% for linear regression and 1.2% for the AMA-based estimator. (d) The contrast normalized cone excitations projected along the first two AMA receptive fields. The receptive fields were learned using AMA and the contrast normalized cone excitations. For Condition 2, the response corresponding to individual image patches at each LRF level separates much better when the input is the contrast normalized cone excitations than when the input is the raw cone excitations.

Recall that there are two qualitatively distinct factors contributing to the variation in illumination spectra: changes in overall intensity, and changes in the relative spectral power distribution. To separate the effect of these two factors, we trained and evaluated AMA on the contrast normalized cone excitations. By contrast normalizing the cone excitations, the contribution of overall intensity is essentially removed. Figure 11(c,d) shows that performance improves greatly with contrast normalization. Estimates obtained from the AMA receptive field responses are essentially perfect. Estimates obtained from linear regression are substantially improved. This observation is consistent with previous results on computational color constancy that show that contrast-based representations are effective for supporting color constancy when only illumination spectra vary (e.g. Land, 1986). However, contrast-based representations are less effective at supporting constancy when the spectra of background objects in the scene vary (e.g. Brainard & Wandell, 1986).

### Condition 3

Condition 3 introduces variation in the reflectance spectra of the background objects, in addition to the sources of variation in Conditions 1 and 2. This condition models the sources of real-world spectral variation that are most relevant for the computational problem of luminance constancy. We again trained and evaluated AMA for labeled image patches, using the contrast normalized cone excitations. The LRF estimates obtained via AMA are more variable and less accurate than for the previous conditions (Figure 12a). Nevertheless, the estimates provide useful information about the target LRF. Indeed, the estimates track the true LRF on average, as seen by the fact that the mean estimate (red line) lies along the positive diagonal. The increased estimation variability is indicated by the increased width of the shaded red region, relative to its width for the results of Condition 1 and 2. Performance of the baseline linear regression method (also evaluated using contrast normalized cone excitations) is worse than that of the AMA-based estimates.

**Figure 12:**
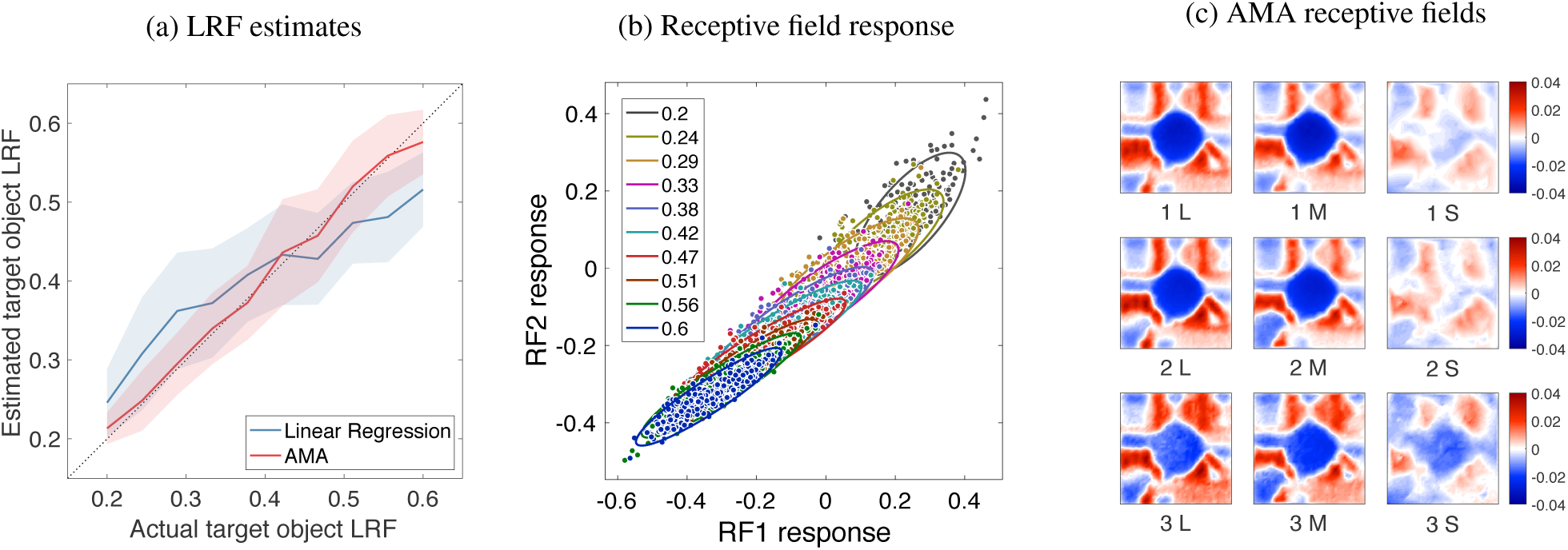
Condition 3 results: (a) Mean estimated LRF ± 1 standard deviation for images in the test set obtained using the AMA-based estimator and linear regression from the contrast normalized cone excitations. The relative RMSE is 23.9% for linear regression and 12.6% for AMA. (b) Contrast normalized cone excitations for the training set projected onto the first two AMA receptive fields. (c) First three AMA receptive fields learned using the contrast normalized cone excitations.

Figure 12b shows the responses of the first two AMA receptive fields to the training stimuli. Although the responses vary systematically with target object LRF, there is considerable overlap in the receptive field responses for stimuli having different LRF values. This overlap is due to the combined effect of the variation in the illumination and background surface spectra.

Recall, however, that the performance shown Figure 12a is based on six rather than two receptive fields; there is likely to be less overlap in the full six-dimensional response space. That is, inclusion of more receptive fields will in general reduce the ambiguity that is seen when we visualize the responses of just the first two receptive fields. This effect is shown in the right panel of Figure 13. That figure also illustrates the effect of training set size on estimation performance.

**Figure 13:**
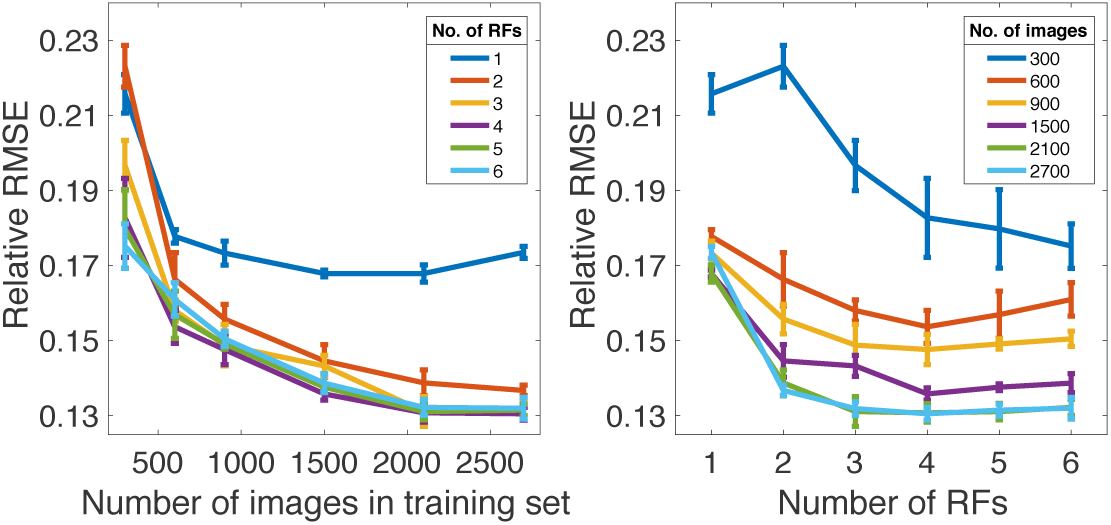
Change of relative RMSE with number of receptive fields (RFs) and size of training set: (a) Change of relative RMSE for Condition 3 with the size of training set. The relative RMSE converges around 2000 images. To make this plot, the 300 image test set for Condition 3 was used in each case, while the training set size varied between 300 to 2700 images. The number of receptive fields used was varied between 1 and 6. (b) Same data as in panel a, but plotted as a function of number of receptive fields. Performance converges around three receptive fields (RFs). The error bars in each panel show +/-1 SEM, with the variation taken across multiple runs with different random initializations for the numerical search used in our implementation of AMA.

Figure 12c shows the first three receptive fields (RFs) learned for Condition 3. The first two are similar to the receptive fields for Condition 1, in that the L and M cone excitations receive large weights, while the S cone excitations are less heavily weighted. In contrast to the Condition 1 receptive fields, there is also systematic contribution of cone excitations from background object locations, with both positive and negative contributions from background objects. The specific geometry of the receptive fields, however, should not be taken to be a strict prediction for receptive fields in the human visual system, as the particular receptive field geometry arises partly as a consequence of the fixed scene geometry used in the training set. Indeed, aspects of spatial structure of the training images can be seen in the receptive fields. Also, the third receptive field for Condition 3 places weight on the S cone excitations. Although S cone signals do not contribute directly to luminance, S cone responses are correlated with L and M cone responses to natural spectra (Burton & Moorhead, 1987; Benson, Manning, & Brainard, 2014). Information from the S cones can therefore contribute usefully to performance in the task. Finally, we note that there is some run-to-run variation in the structure of the optimal receptive fields that depends on the random initialization of the numerical search used in our implementation of AMA. (The cost landscape is not convex so local minima are possible.) However, such run-to-run receptive field variation leads only to very small changes in estimation error (see error bars in Figure 13). Furthermore, even if the order varies somewhat, the same set of receptive fields tends to be learned.

Our rendering software allows us to compare the effect of background surface reflectances on target object LRF with and without simulation of secondary reflections of light from one object onto another. These secondary reflections were included in the dataset from which we report our primary results. When we turn off this feature of the rendering, we find (data not shown) that LRF estimation performance is essentially unchanged. Estimates with and without secondary reflections are very similar. This result suggests that the primary source of the estimation error in Condition 3 is caused by direct effects of image-to-image variation in the reflectance of the background objects on the AMA responses.

## Discussion

### Summary

In this paper, we studied luminance constancy using naturalistic images. We used a software pipeline, which was developed for this work, to render datasets of multispectral images from scene descriptions. Because we rendered the images, we could label each image in the dataset by the luminous reflectance factor (LRF) of the target object. We used the labeled datasets to learn estimators for the target object LRF. Across scenes, we varied the LRF of the target object, the relative reflectance spectrum of the target object, the spectral power distributions of the light sources, and the reflectance spectra of the background objects in the scene. These spectral variations were based on statistical models of natural surface reflectance spectra and natural illumination spectra. We studied the how the performance of the learned estimators changed with systematic manipulation of the spectral variation in the datasets.

Figure 14 summarizes our main findings, by showing the overall relative root mean squared estimation error (relative RMSE) for each condition. In Condition 1, where only the relative reflectance spectrum of the target object varies, luminance constancy is an easy computational problem, and performance is essentially perfect for both the baseline linear regression and AMA methods. In Condition 2, where the spectral power distribution of the illumination also varies, the problem of luminance constancy is more difficult. Indeed, neither of our methods perform well when they operate directly on the cone excitations. Performance with both methods improves greatly when the cone excitations are contrast normalized. Indeed, AMA performance for Condition 2 is essentially perfect, as it is in Condition 1. Linear regression does less well, in part because it only has access to cone excitations at image locations corresponding to the target object. Condition 2, however, does not include scene-to-scene variation in the background surface reflectances, as occurs in natural viewing. The results for Condition 3 show performance when we add such variation. Performance here represents the overall level of luminance constancy that our methods achieve for the most realistic condition that we tested. In this case, AMA-based estimates of target object LRF are, on average, within 13% (relative RMSE) of the true value. Introducing variation in the surface reflectance spectra of the background objects makes the luminance constancy problem substantially more difficult. Variation in the reflectance spectra of the background objects surfaces reduces how reliably light reflected from those objects can be used as a cue to estimate the spectral power distribution of the illuminant.

**Figure 14:**
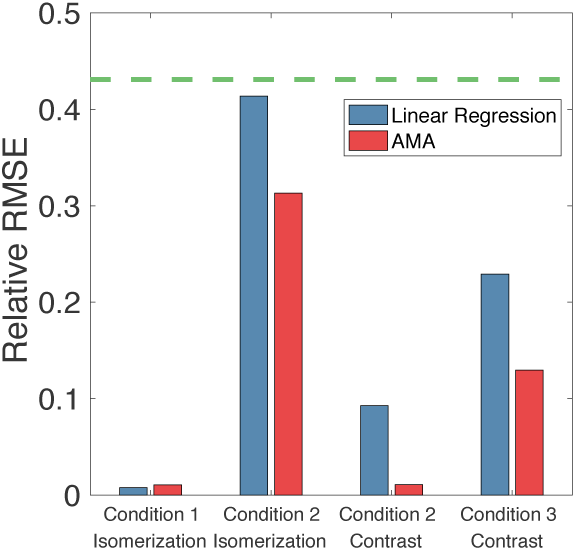
Summary of luminance constancy performance: Relative RMSE estimation error for each method and condition. The dotted line represents the relative RMSE for a naive method that estimates the LRF of each image as the mean LRF over the images in the set (LRF = 0.4).

### Images of Virtual vs. Real Scenes

As noted in the introduction, supervised learning requires large labeled datasets. Labeled datasets of natural images have been useful for developing normative models of the estimation of defocus blur, binocular disparity, retinal image motion, and 3D surface orientation (Burge & Geisler, 2011, 2012, 2014, 2015; Sebastian, Burge, & Geisler, 2015; Kim & Burge, 2018; Burge, Fowlkes, & Banks, 2010; Girshick, Landy, & Simoncelli, 2011; Burge, McCann, & Geisler, 2016; Goncalves & Welchman, 2017).

Large datasets of natural or posed scenes with ground truth information about illuminant and object surface reflectance are difficult to obtain, as independent measurement of illuminant and surface properties at each image location is painstaking. That noted, there are databases for evaluation of color constancy algorithms that provide information about illumination and/or surface reflectance (e.g., Barnard, Martin, Funt, & Coath, 2002; Ciurea & Funt, 2003; Cheng, Prasad, & Brown, 2014; Nascimento et al., 2016; see http://colorconstancy.com). Often the illumination is estimated through placement of a reflectance standard at a few image locations to allow estimation of the illumination impinging at those locations. These illumination estimates are then interpolated/extrapolated across the image. However, the quality of this approximation cannot typically be evaluated.

Here we used labeled images rendered from descriptions of virtual scenes. A similar approach has been used previously to study the perception of lightness and specularity (Toscani, Valsecchi, & Gegenfurtner, 2013; Wiebel, Toscani, & Gegenfurtner, 2015; Toscani, Valsecchi, & Gegenfurtner, 2017). Our work adds to this approach by introducing color variation. There are many advantages to using rendered images (Butler, Wulff, Stanley, & Black, 2012). One advantage is that they allow us to work with large number of labeled images where object reflectance is precisely known at each pixel. A second advantage is that we can control the variation in distinct scene factors that might affect the difficulty of the estimation problem. This flexibility allows the impact of scene factors to be studied individually or in combination. Here, we exploited this flexibility to quantify how variation in the relative reflectance spectrum of the target object, the spectrum of the illumination, and the reflectance spectrum of the background objects limit LRF estimation. We also exploited our use of rendered images to explore how the presence or absence of secondary reflections from background objects affected estimation of target object LRF. This type of question cannot be addressed using real images. The basic approach we use here can be extended to include parametric control over the amount of variation of different factors. For example, we could systematically vary the variances of the distribution over the weights that control the relative spectrum of the illumination.

There are also disadvantages associated with using rendered images. Virtual rendered scenes are not guaranteed to capture all of the task relevant variation that occurs in real scenes. Even casual inspection of our images reveals that they are rendered, and not real. However, computer graphics is getting better and we expect that the gap between virtual and natural image databases will steadily close. Indeed, carefully constructed graphics images are now quite difficult to differentiate from real images.

To increase the representativeness of our rendered images, we used datasets of natural surface reflectance spectra and natural daylight illumination spectra. Although we believe these datasets provide reasonable approximations of the statistical variation in reflectance and illumination spectra, they can be extended and improved. For example, there are additional datasets of measured surface reflectances that could be incorporated into future analyses. Some of these datasets focused on the reflectance of objects (e.g. fruit) that are thought to be important for the evolution of primate color vision (e.g., Sumner & Mollon, 2000; Regan et al., 2001; Barnard et al., 2002; Ennis, Schiller, Toscani, & Gegenfurtner, 2018). Another issue, not addressed by these datasets, is relative frequency of different surface reflectances in natural viewing. Attewell & Baddeley, 2007 performed a systematic survey, and reported the distribution of an LRF-like quantity in natural scenes. Generalizing these measurements to better characterize the distribution of full reflectance functions remains an interesting goal.

### Future Directions

In the work presented here, we studied computational luminance constancy in virtual scenes with naturalistic spectral variation in light sources and in surface reflectance functions, with only matte surfaces in the scenes. It is natural to start with spectral variation, because this variation is at the heart of what makes luminance constancy a rich computational problem. In natural scenes, however, there are other sources of variation that add additional richness. These include variation in non-spectral properties of lighting and objects in the scene. Examples include lighting geometry, object texture, object material (e.g. specularity), and object shape. The methods we developed here may be generalized to study the effects of variation in these factors. That is, one could incorporate these factors into the generation of the scenes and again learn estimators from the corresponding labeled images. A challenge for this approach will be to thoughtfully control the increase in problem complexity, both to keep compute time feasible and to ensure that it is possible to extract meaningful insight from the results. Extending the work to include variation of material may provide insights not only about luminance constancy but also for computations that relate to material perception (see Fleming, 2017). Extending the work to include variation in object shape and lighting geometry may clarify the role of object boundaries versus object interiors for providing information that supports perception of object color and lightness (see Land & McCann, 1971; Rudd, 2016). We also note that there is a literature on how increasing stimulus complexity along the various lines listed above affects human color and lightness perception (e.g. Beck, 1964; Yang & Maloney, 2001; Yang & Shevell, 2002; Todd, Norman, & Mingolla, 2004; Snyder, Doerschner, & Maloney, 2005; Boyaci, Maloney, & Hersh, 2003; Xiao & Brainard, 2008; Kingdom, 2011; Xiao, Hurst, MacIntyre, & Brainard, 2012; Anderson, 2015; Toscani et al., 2017), and the computational problem of color and lightness constancy (e.g. Lee, 1986; D’Zmura & Lennie, 1986; Funt & Drew, 1988; Tominaga & Wandell, 1989; Barron & Malik, 2012; Barron, 2015; Finlayson, 2018).

We studied the information available for LRF estimation using a 1° × 1° image patch. As noted above this choice of size was motivated in part to use a spatial scale roughly commensurate with the scale of information integration in early visual cortex. Our general methods could be extended to study larger regions, and doing so would quantify the value of spatially remote information for luminance constancy.

Luminance constancy is a special case of the more general problem of color constancy. The approach we developed here could also be generalized to the estimation of additional surface reflectance descriptors, such as object hue or chroma. A further advance would be to develop methods for learning receptive fields that are optimal for multivariate estimation problems. With such methods one could directly estimate three-dimensional color descriptors.

### Linking to Human Performance

An important motivation for studying the computational problem of luminance constancy is to gain insight about human vision. The approach we developed can be used to make predictions of how humans would perform in a psychophysical task that probes the visual representation of object LRF. For example, one could study discrimination thresholds for target LRF using the stimuli that were generated with the methods used here. More specifically, we could ask how LRF discrimination thresholds are impacted by spectral variation in the illuminant, the background objects, and in the target object itself. Comparing human thresholds with the precision of AMA-based estimates would then allow inferences about how well human observers make use of the information available in the images for performing LRF discrimination. We think experiments and analyses along these lines represent an important future direction.

## Acknowledgments

Supported by startup funds from the University of Pennsylvania (JB), NIH grant R01 EY028571 from the National Eye Institute and the Office of Behavioral and Social Science Research (JB), NIH grant RO1 EY100106 (DHB) from the National Eye Institute, Simons Foundation Collaboration on the Global Brain Grant 324759 (DHB), and the University of Pennsylvania Computational Neuroscience Initiative (VS). Preliminary versions of this work were presented at the Computational Cognitive Neuroscience Conference (2017) and the Annual Meeting of the Society for Neuroscience (2017).

## Appendix

### 1 Illumination Spectra

Denote the *i*’th spectrum in the Granada dataset as 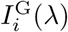, where {*i ∈* [1*,M*]} and *M* is the total number of spectra in the dataset. Since the measured spectra vary over several orders of magnitude in overall intensity, we normalize each spectrum by dividing it by its mean power to obtain what we refer to as rescaled spectra:

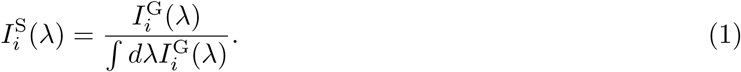

For simplicity of notation, we denote wavelength λ as a continuous variable; in the actual calculations wavelength is discretely sampled and integrals are approximated by sums. The Granada dataset was measured at 5 nm sampling intervals between 300 and 1100 nm. We subsampled the spectra to the 400-700 nm interval, 10 nm spacing representation used for rendering, and performed our calculations at this sampling.

The rescaled spectra 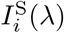 were mean centered for PCA by subtracting out the mean 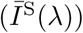, over all rescaled spectra in the dataset,

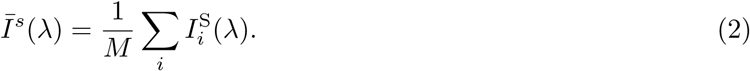

The mean centered dataset, 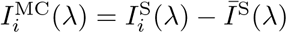 was decomposed as:

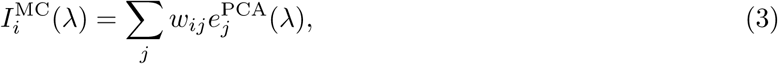

where the 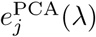 are the PCA basis vectors obtained using singular value decomposition (SVD) applied to 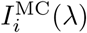 and *w*_*ij*_ are the weights obtained by projecting each of 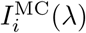 onto the basis vectors 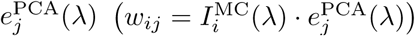 We used the basis vectors corresponding to the largest six SVD eigenvalues, so that {*j∈* [1, 6]. For the rescaled Granada dataset, these six eigenvalues account for more than 90% of the variance.

To get a random illuminant spectrum 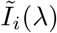, we sample random weights 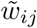 from a multivariate Gaussian distribution with mean 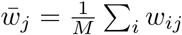, and covariance matrix Σ given as:

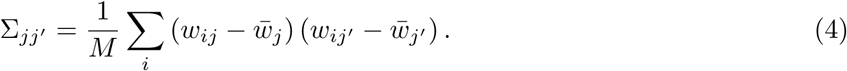

The corresponding spectrum is generated from these weights as 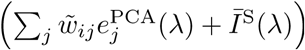 This spectrum can sometimes have values that are less than zero. In such cases, the weights are discarded and a new draw obtained, until the condition 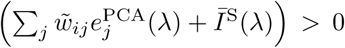, is satisfied for all λ. The random illuminant spectrum is

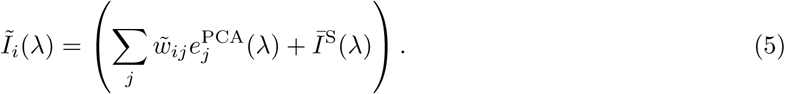

#### 2 Surface Reflectance Spectra

Denote the *i*’th sample in the reflectance spectrum dataset as *R*_*i*_ (λ), where {*i* ∈ [1*,M*]} and *M* is the total number of spectra in the dataset. We first calculated the mean reflectance spectrum, 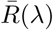, by taking the sample mean over all spectra in the dataset

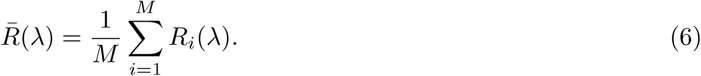

The dataset is then mean centered by subtracting the mean spectrum, 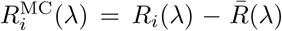 and decomposed using PCA as:

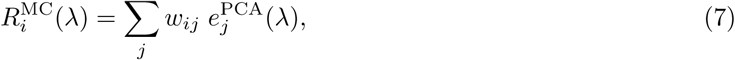

where 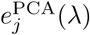 are PCA basis vectors obtained using SVD applied to 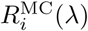 and *w*_*ij*_ are the weights obtained by projecting each of 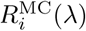 onto the basis vectors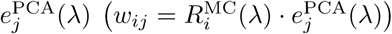. We used the basis vectors corresponding to the largest six SVD eigenvalues, which account for more than 90% of the variance in the combined Munsell and Vrhel datasets.

To get a random reflectance spectrum, we generate samples of weights (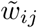) drawn from the multivariate Gaussian distribution with mean 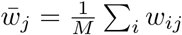, and covariance 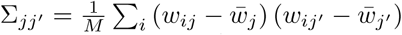. If the randomly sampled weights satisfy the condition 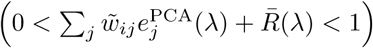 at every λ randomly generated reflectance spectrum is given as:

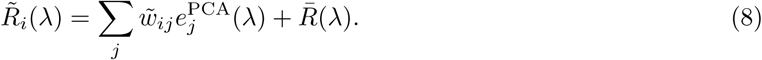

Otherwise the draw is discarded and a new set of weights is drawn.

For generating the target object reflectance at a particular target LRV (*Y*_T_), the values in a generated spectrum were scaled by

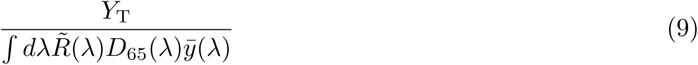

with 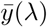 being the CIE photopic luminosity (or luminous efficiency) function, which describes the average spectral sensitivity of human visual perception of luminance. The target reflectance is then given by:

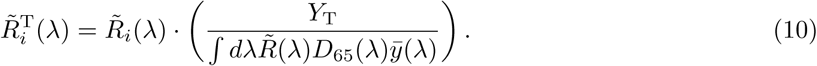

**Figure 1:**
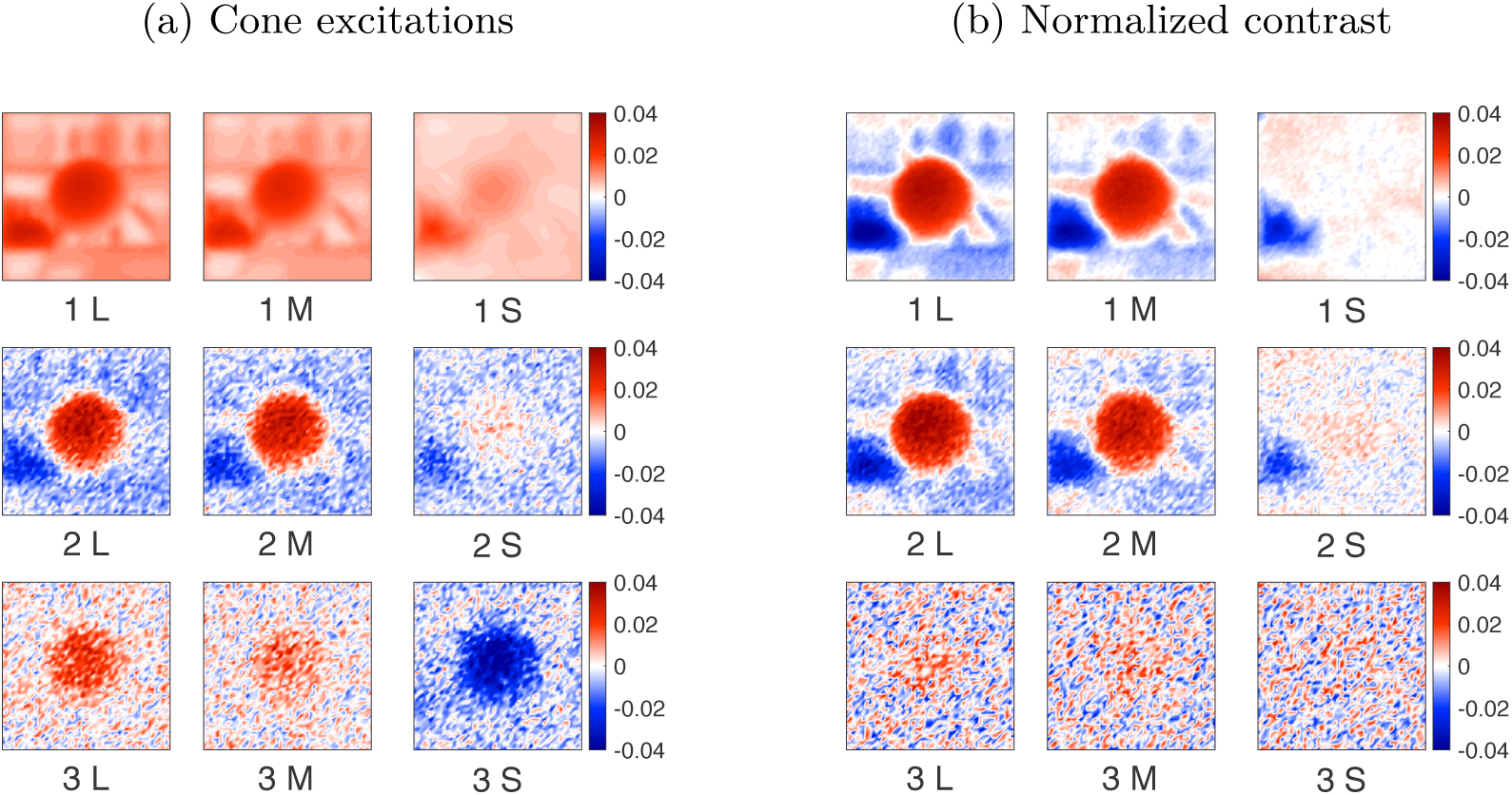
Condition 2 receptive fields: The first three AMA receptive fields learnt on (a) Cone excitations (b) Normalized contrast.

## Footnotes

^1^In the literature, the term lightness constancy is generally used to denote color constancy for the special case when stimuli are restricted to be achromatic. This condition does not apply to our work - we consider full spectral variation in the stimuli. We chose the term luminance constancy to denote the generalization from achromatic stimuli. At the same time we acknowledge that we are not studying the full problem of color constancy. Rather, we are studying the estimation of a luminance-based summary of surface spectral reflectance.

^2^LRF is related to albedo, but the concept of albedo does not incorporate the human luminosity function.

## References

Adelson, E. H. (2000). Lightness perception and lightness illusions. In M. Gazzaniga (Ed.), The New Cognitive Neurosciences. Cambridge, MA: MIT Press, 339–351.

Albrecht, D. G., & Geisler, W. S. (1991). Motion selectivity and the contrast-response function of simple cells in the visual cortex. Visual Neuroscience, 7(6), 531–546.

American Society for Testing and Materials. (2017). Standard test method for luminous reflectance factor of acoustical materials by use of integrating-sphere reflectometers. Renovations of Center for Historic Preservation, 98(A), E1477.

Anderson, B. L. (2015). The perceptual representation of transparency, lightness, and gloss. In J. Wagemans (Ed.), Handbook of Perceptual Organization. Oxford: Oxford University Press.

Attewell, D., & Baddeley, R. J. (2007). The distribution of reflectances within the visual environment. Vision Research, 47(4), 548–554.

Barnard, K., Martin, L., Funt, B., & Coath, A. (2002). A data set for color research. Color Research & Application, 27(3), 147–151.

Barron, J. T. (2015). Convolutional color constancy. Proceedings of the IEEE International Conference on Computer Vision, 379–387.

Barron, J. T., & Malik, J. (2012). Color constancy, intrinsic images, and shape estimation. Proceedings of the European Conference on Computer Vision (ECCV), 57–70.

Beck, J. (1964). The effect of gloss on perceived lightness. The American Journal of Psychology, 77(1), 54–63.

Bell, S., Bala, K., & Snavely, N. (2014). Intrinsic images in the wild. ACM Transactions on Graphics (TOG), 33(4), 159.

Benson, N. C., Manning, J. R., & Brainard, D. H. (2014). Unsupervised learning of cone spectral classes from natural images. PLoS Computational Biology, 10(6), e1003652.

Boyaci, H., Maloney, L. T., & Hersh, S. (2003). The effect of perceived surface orientation on perceived surface albedo in binocularly viewed scenes. Journal of Vision, 3(8), 541–553.

Brainard, D. H. (2015). Color and the cone mosaic. Annual Review of Vision Science, 1, 519–546.

Brainard, D. H., & Freeman, W. T. (1997). Bayesian color constancy. Journal of the Optical Society of America A, 14(7), 1393–1411.

Brainard, D. H., & Radonjić, A. (2014). Color constancy. In L. M. Chalupa & J. S. Werner (Eds.), The New Visual Neurosciences. MIT Press, 545–556.

Brainard, D. H., & Wandell, B. A. (1986). Analysis of the retinex theory of color vision. Journal of the Optical Society of America A, 3(10), 1651–1661.

Buchsbaum, G. (1980). A spatial processor model for object colour perception. Journal of the Franklin Institute, 310(1), 1–26.

Burge, J., Fowlkes, C. C., & Banks, M. S. (2010). Natural-scene statistics predict how the figure–ground cue of convexity affects human depth perception. Journal of Neuroscience, 30(21), 7269–7280.

Burge, J., & Geisler, W. S. (2011). Optimal defocus estimation in individual natural images. Proceedings of the National Academy of Sciences, 108(40), 16849–16854.

Burge, J., & Geisler, W. S. (2012). Optimal defocus estimates from individual images for autofocusing a digital camera. Digital Photography VIII, 8299, 82990E.

Burge, J., & Geisler, W. S. (2014). Optimal disparity estimation in natural stereo images. Journal of Vision, 14(2), 1–18.

Burge, J., & Geisler, W. S. (2015). Optimal speed estimation in natural image movies predicts human performance. Nature Communications, 6, 7900.

Burge, J., & Jaini, P. (2017). Accuracy maximization analysis for sensory-perceptual tasks: Computational improvements, filter robustness, and coding advantages for scaled additive noise. PLoS Computational Biology, 13(2), e1005281.

Burge, J., McCann, B. C., & Geisler, W. S. (2016). Estimating 3d tilt from local image cues in natural scenes. Journal of Vision, 16(13), 1–25.

Burton, G., & Moorhead, I. R. (1987). Color and spatial structure in natural scenes. Applied Optics, 26(1), 157–170.

Butler, D. J., Wulff, J., Stanley, G. B., & Black, M. J. (2012). A naturalistic open source movie for optical flow evaluation. European Conference on Computer Vision (ECCV), 611–625.

Carandini, M., & Heeger, D. J. (2012). Normalization as a canonical neural computation. Nature Reviews Neuroscience, 13(1), 51–62.

Chakrabarti, A., & Zickler, T. (2011). Statistics of real-world hyperspectral images. Proceedings of the IEEE Computer Society Conference on Computer Vision and Pattern Recognition, 193–200.

Cheng, D., Prasad, D. K., & Brown, M. S. (2014). Illuminant estimation for color constancy: why spatial-domain methods work and the role of the color distribution. Journal of the Optical Society of America A, 31(5), 1049–1058.

Ciurea, F., & Funt, B. (2003). A large image database for color constancy research. Color and Imaging Conference, 2003(1), 160–164.

Commission Internationale de l’éclairage. (1986). Colorimetry, second edition (Tech. Rep. No. 15.2). Bureau Central de la CIE.

D’Zmura, M., & Iverson, G. (1994). Color constancy. iii. General linear recovery of spectral descriptions for lights and surfaces. Journal of the Optical Society of America A, 11(9), 2389–2400.

D’Zmura, M., Iverson, G., & Singer, B. (1995). Probabilistic color constancy. Geometric Representations of Perceptual Phenomena: Papers in Honor of Tarow Indow’s 70th Birthday, 187–202.

D’Zmura, M., & Lennie, P. (1986). Mechanisms of color constancy. Journal of the Optical Society of America A, 3(10), 1662–1672.

Ennis, R., Schiller, F., Toscani, M., & Gegenfurtner, K. R. (2018). Hyperspectral database of fruits and vegetables. Journal of the Optical Society of America A, 35(4), B256–B266.

Finlayson, G. D. (2018). Colour and illumination in computer vision. Interface Focus, 8(4), 20180008.

Fleming, R. W. (2017). Material perception. Annual Review of Vision Science, 3, 365–388.

Foster, D. H. (2011). Color constancy. Vision Research, 51(7), 674–700.

Funt, B. V., & Drew, M. S. (1988). Color constancy computation in near-mondrian scenes using a finite dimensional linear model. Computer Society Conference on Computer Vision and Pattern Recognition, 544–549.

Gattass, R., Gross, C., & Sandell, J. (1981). Visual topography of V2 in the macaque. Journal of Comparative Neurology, 201(4), 519–539.

Gattass, R., Sousa, A., & Gross, C. (1988). Visuotopic organization and extent of V3 and V4 of the macaque. Journal of Neuroscience, 8(6), 1831–1845.

Geisler, W. S., Najemnik, J., & Ing, A. D. (2009). Optimal stimulus encoders for natural tasks. Journal of Vision, 9(13), 1–6.

Gilchrist, A. (2006). Seeing black and white. Oxford: Oxford University Press.

Girshick, A. R., Landy, M. S., & Simoncelli, E. P. (2011). Cardinal rules: visual orientation perception reflects knowledge of environmental statistics. Nature Neuroscience, 14(7), 926–932.

Goncalves, N. R., & Welchman, A. E. (2017). “What not” detectors help the brain see in depth. Current Biology, 27(10), 1403–1412.

Heasly, B. S., Cottaris, N. P., Lichtman, D. P., Xiao, B., & Brainard, D. H. (2014). Rendertoolbox3: Matlab tools that facilitate physically based stimulus rendering for vision research. Journal of Vision, 14(2), 1–22.

Hecht, S., Shlaer, S., & Pirenne, M. H. (1942). Energy, quanta, and vision. The Journal of General Physiology, 25(6), 819–840.

Heeger, D. J. (1992). Normalization of cell responses in cat striate cortex. Visual Neuroscience, 9(2), 181–197.

Hernández-Andrés, J., Romero, J., Nieves, J. L., & Lee, R. L. (2001). Color and spectral analysis of daylight in southern europe. Journal of the Optical Society of America A, 18(6), 1325–1335.

Jacobs, G. H. (1981). Comparative Color Vision. New York: Academic Press.

Jaini, P., & Burge, J. (2017). Linking normative models of natural tasks to descriptive models of neural response. Journal of Vision, 17(12), 1–26.

Jakob, W. (2010). Mitsuba renderer. http://www.mitsuba-renderer.org.

Kelly, K. L., Gibson, K. S., & Nickerson, D. (1943). Tristimulus specification of the Munsell book of color from spectrophotometric measurements. Journal of the Optical Society of America, 33(7), 355–376.

Kim, S., & Burge, J. (2018). The lawful imprecision of human surface tilt estimation in natural scenes. eLife, 7, e31448.

Kingdom, F. A. (2011). Lightness, brightness and transparency: A quarter century of new ideas, captivating demonstrations and unrelenting controversy. Vision Research, 51(7), 652–673.

Land, E. H. (1977). The retinex theory of color vision. Scientific American, 237(6), 108–128.

Land, E. H. (1986). An alternative technique for the computation of the designator in the retinex theory of color vision. Proceedings of the National Academy of Sciences, 83(10), 3078–3080.

Land, E. H., & McCann, J. J. (1971). Lightness and retinex theory. Journal of the Optical Society of America, 61(1), 1–11.

Lee, H.-C. (1986). Method for computing the scene-illuminant chromaticity from specular highlights. Journal of the Optical Society of America A, 3(10), 1694–1699.

Maloney, L., & Wandell, B. A. (1986). Color constancy: a method for recovering surface spectral reflectances. Journal of the Optical Society of America A, 3, 29–33.

Marimont, D. H., & Wandell, B. A. (1994). Matching color images: the effects of axial chromatic aberration. Journal of the Optical Society of America A, 11(12), 3113–3122.

Mollon, J. D. (1989). “Tho’ she kneel’d in that place where they grew…” The uses and origins of primate colour vision. Journal of Experimental Biology, 146, 21–38.

Nascimento, S. M., Amano, K., & Foster, D. H. (2016). Spatial distributions of local illumination color in natural scenes. Vision Research, 120, 39–44.

Olmos, A., & Kingdom, F. A. (2004). A biologically inspired algorithm for the recovery of shading and reflectance images. Perception, 33(12), 1463–1473.

Parraga, C. A., Brelstaff, G., Troscianko, T., & Moorehead, I. R. (1998). Color and luminance information in natural scenes. Journal of the Optical Society of America A, 15, 563–569.

Regan, B. C., Julliot, C., Simmen, B., Vienot, F., Charles-Dominique, P., & Mollon, J. D. (2001). Fruits, foliage and the evolution of primate colour vision. Philosophical Transactions of the Royal Society of London B: Biological Sciences, 356(1407), 229–283.

Rudd, M. E. (2016). Retinex-like computations in human lightness perception and their possible realization in visual cortex. Electronic Imaging, 2016(6), 1–8.

Sebastian, S., Burge, J., & Geisler, W. S. (2015). Defocus blur discrimination in natural images with natural optics. Journal of Vision, 15(5).

Skauli, T., & Farrell, J. (2013). A collection of hyperspectral images for imaging systems research. Digital Photography IX, 8660, 86600C1–C7.

Snyder, J. L., Doerschner, K., & Maloney, L. T. (2005). Illumination estimation in three-dimensional scenes with and without specular cues. Journal of Vision, 5(10), 863–877.

Sumner, P., & Mollon, J. D. (2000). Catarrhine photopigments are optimized for detecting targets against a foliage background. Journal of Experimental Biology, 203(13), 1963–1986.

Tkacik, G., Garrigan, P., Ratliff, C., Milcinski, G., Klein, J. M., Sterling, P., Brainard, D. H., & Balasubramanian, V. (2011). Natural images from the birthplace of the human eye. PLoS ONE, 6(6), e20409.

Todd, J. T., Norman, J. F., & Mingolla, E. (2004). Lightness constancy in the presence of specular highlights. Psychological Science, 15(1), 33–39.

Tominaga, S., & Wandell, B. A. (1989). Standard surface-reflectance model and illuminant estimation. Journal of the Optical Society of America A, 6(4), 576–584.

Toscani, M., Valsecchi, M., & Gegenfurtner, K. R. (2013). Optimal sampling of visual information for lightness judgments. Proceedings of the National Academy of Sciences, 110(27), 11163–11168.

Toscani, M., Valsecchi, M., & Gegenfurtner, K. R. (2017). Lightness perception for matte and glossy complex shapes. Vision Research, 131, 82–95.

Vrhel, M. J., Gershon, R., & Iwan, L. S. (1994). Measurement and analysis of object reflectance spectra. Color Research & Application, 19(1), 4–9.

Wiebel, C. B., Toscani, M., & Gegenfurtner, K. R. (2015). Statistical correlates of perceived gloss in natural images. Vision Research, 115, 175–187.

Xiao, B., & Brainard, D. H. (2008). Surface gloss and color perception of 3d objects. Visual Neuroscience, 25(3), 371–385.

Xiao, B., Hurst, B., MacIntyre, L., & Brainard, D. H. (2012). The color constancy of three-dimensional objects. Journal of Vision, 12(4), 6–6.

Yang, J. N., & Maloney, L. T. (2001). Illuminant cues in surface color perception: Tests of three candidate cues. Vision Research, 41(20), 2581–2600.

Yang, J. N., & Shevell, S. K. (2002). Stereo disparity improves color constancy. Vision Research, 42(16), 1979–1989.

